# Fluid Flow Impacts Endothelial-Monocyte Interactions in a Model of Vascular Inflammatory Fibrosis

**DOI:** 10.1101/2024.09.29.615690

**Authors:** Isabelle Linares, Kaihua Chen, Ava Saffren, Mehran Mansouri, Vinay V. Abhyankar, Benjamin L. Miller, Stefano Begolo, Hani A. Awad, James L. McGrath

## Abstract

The aberrant vascular response associated with tendon injury results in circulating immune cell infiltration and a chronic inflammatory feedback loop leading to poor healing outcomes. Studying this dysregulated tendon repair response in human pathophysiology has been historically challenging due to the reliance on animal models. To address this, our group developed the human tendon-on-a-chip (hToC) to model cellular interactions in the injured tendon microenvironment; however, this model lacked the key element of physiological flow in the vascular compartment. Here, we leveraged the modularity of our platform to create a fluidic hToC that enables the study of circulating immune cell and vascular crosstalk in a tendon injury model. Under physiological shear stress consistent with postcapillary venules, we found a significant increase in the endothelial leukocyte activation marker intercellular adhesion molecule 1 (ICAM-1), as well as enhanced adhesion and transmigration of circulating monocytes across the endothelial barrier. The addition of tissue macrophages to the tendon compartment further increased the degree of circulating monocyte infiltration into the tissue matrix. Our findings demonstrate the importance of adding physiological flow to the human tendon-on-a-chip, and more generally, the significance of flow for modeling immune cell interactions in tissue inflammation and disease.

## Introduction

Tendons are a vital component of the human musculoskeletal system, connecting muscle to bone and transducing forces to produce movement.^[1]^ Due to overuse and age-related degeneration, tendon injuries have become an increasingly significant clinical problem. Approximately 32 million cases of tendon injury are reported in the United States each year, accounting for 30% of musculoskeletal consultations and producing an estimated $30 billion economic burden due to healthcare expenses and loss of productivity.^[2–4]^

Healthy tendons are hypovascular with a relatively low population of cells and only 1%-2% of the collagen matrix occupied by vessels.^[5]^ In contrast, injured tendons are characterized by chronic inflammation leading to a vascularized fibrotic scar and impaired mechanical properties.^[6]^ An increase in vasculature following tendon injury is associated with degeneration and is a key factor in promoting a chronic fibrotic response.^[7]^ This dysregulated healing response is, in part, attributed to the increased activation of myofibroblasts by secreted cytokines released by tissue-resident immune cells, endothelial cells (ECs), and infiltrating leukocytes including neutrophils and monocytes.

The extent of neutrophil and monocyte infiltration following injury plays a key role in determining the progression of tendon repair toward scarring.^[8]^ Within 2–3 days following injury, pro-inflammatory cytokine levels at the defect site increase several thousand times and drive the transendothelial migration of circulating leukocytes.^[9]^ Among others, tumor necrosis factor-alpha (TNF-α), interleukin-1 beta (IL-1β), transforming growth factor-beta 1 (TGF-β1), and monocyte chemoattractant protein-1 (MCP-1) are primary cytokines upregulated in the post-injury microenvironment.^[3,10]^ Tendon fibroblasts secrete a collagenous matrix containing chemotactic components, which further stimulates fibroblast migration, proliferation, and a transition to a myofibroblast phenotype.^[11]^ However, the underlying mechanism of dysregulated tendon repair and the role of vascular cell types remains unclear. The need to elucidate the effects of infiltrating vasculature and immune cells has motivated the development of physiologically relevant tendon injury models.^[10,12]^

The majority of musculoskeletal models to date are high-cost, low-throughput animal models or over-simplified 2D *in vitro* models, both with limited translational value.^[13]^ To enable the reductionist studies needed to understand the complex physiology of tissue microenvironments, microphysiological systems (MPS), or ‘tissue chips’, have emerged as paradigm-shifting technologies that mimic the essential features of human tissue *in vitro*. To begin addressing the debilitating impact of fibrosis following tendon injury, our laboratory developed a human tendon-on-a-chip (hToC) that features a 3D tendon model and a vascular compartment separated by a highly permeable and ultrathin nanomembrane to enable facile studies of cellular and molecular crosstalk between ‘blood’ and ‘tissue’.^[10]^ While able to recapitulate many of the salient features of injured tendon, the original hToC model is missing the important physiological elements of fluid flow and shear stress at the vascular wall. Both of these strongly impact endothelial cell morphology and function. The range of shear stress on the endothelium varies between 1– 30 dynes/cm^2^ throughout the vascular network.^[14]^ At the postcapillary venule, where immune cell extravasation is most prominent, shear stress falls within 1.5–4.5 dynes/cm^2^.^[15]^ Shear stress ≥10 dynes/cm^2^ promotes endothelial cell elongation and cytoskeletal alignment and improves barrier function.^[16–19]^ Vascular flow also plays a vital role in the clearance of secreted inflammatory mediators and facilitates immune cell infiltration following the physiological cascade that involves rolling, arrest, and transmigration.^[20,21]^ Thus, we hypothesize that fluid flow will modulate the vascular inflammatory response in the hToC model by impacting endothelial barrier function, surface adhesion protein expression, and the rate of immune cell infiltration.

To investigate this hypothesis, we developed a ‘manufacturable’ plug-and-play module that transforms the static hToC into a fluidic hToC enabling improved mechanistic studies of the vasculature. We validated, both computationally and experimentally, that the flow insert replicates physiological shear stress values and preserves characteristic endothelial morphology under applied shear stress. Importantly, we demonstrated that the endothelium exhibits distinct responses to inflammatory stimuli in static versus fluidic culture conditions and that vascular flow augments monocyte adhesion and transmigration. In the context of the complete hToC model, which includes a tenocyte-laden tissue compartment, we optimized a fluidic culture protocol and highlighted the importance of tissue macrophage-derived signaling in driving circulating monocyte infiltration. By incorporating the essential vascular cue of fluid flow, the fluidic hToC allows us to investigate the role of the vascular inflammatory response in fibrotic tendon pathology.

## Results

### Development of a Peel-and-Stick, Manufactured Flow Insert for the hToC

Our laboratory has developed a modular device platform for barrier tissue modeling with ultrathin porous silicon membranes termed the µSiM.^[22]^ The hToC is an extension of the µSiM that replaces the bottom component with a thicker channel to model a 3D matrix tissue. A key advantage of our modular architecture is that the membrane and device components are commercially manufactured at scale, allowing the assembly of individual devices on demand in the laboratory. Assembly takes only minutes and involves activating pressure-sensitive adhesives (PSAs) by removing protective layers and pressing parts together in assembly jigs.^[22]^ While we have previously developed and characterized a hand-crafted flow module for the µSiM^[20,23]^, here we sought to extend the scaled manufacturing and ‘peel-and-stick’ functionality of the modular µSiM to the fluidic hToC. We developed a simple and manufacturable flow module that: 1) creates an aligned flow channel over the membrane with flanking inlet and outlet ports and 2) includes a protected pressure-sensitive adhesive (PSA) layer at the bottom of the channel that can be exposed and sealed against the membrane chip **(Fig. 1)**.

**Figure 1.**
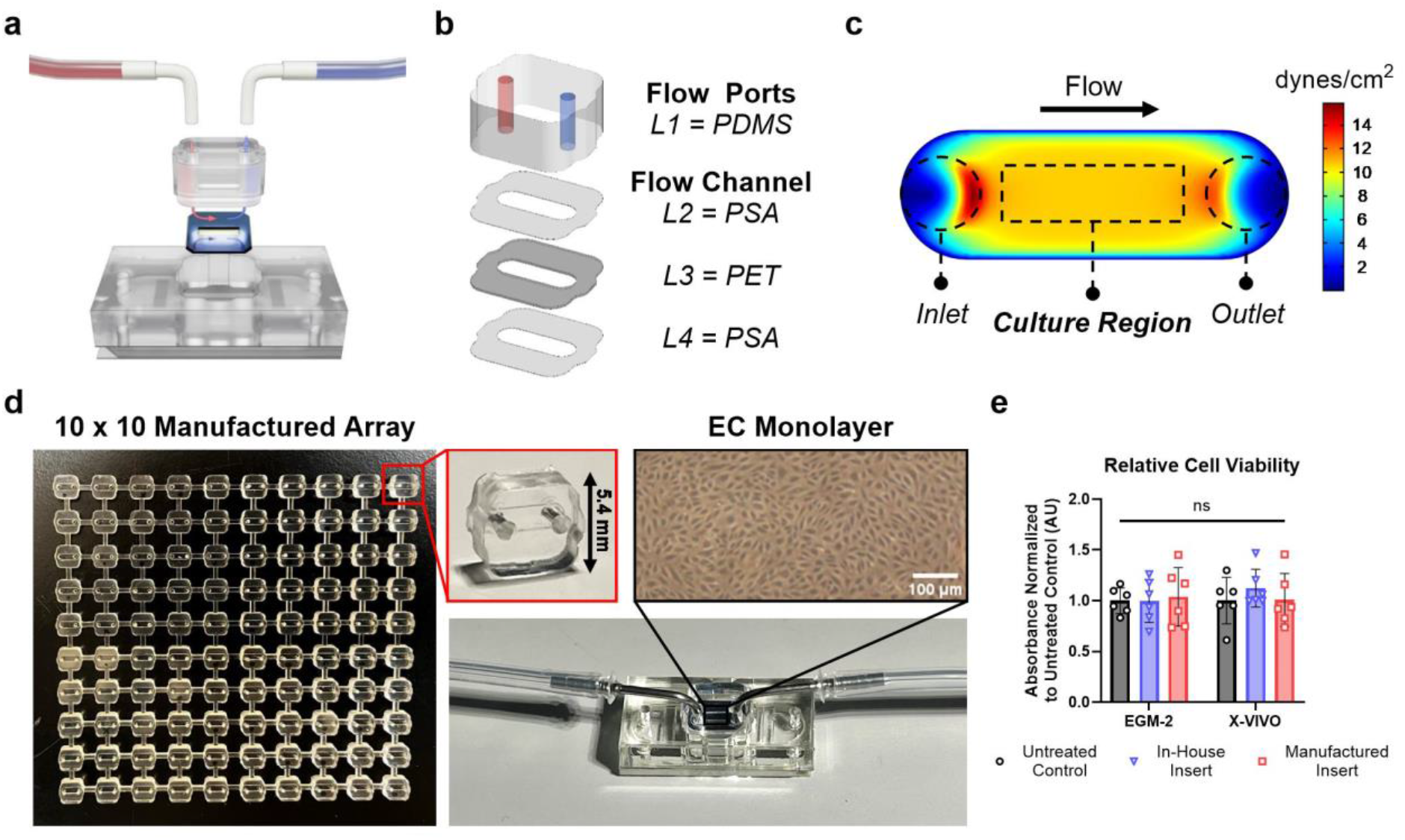
Manufactured flow insert development and validation. **a.** Schematic of fluidic hToC device with flow insert and tubing, indicating inlet flow in red and outlet flow in blue across the ultrathin, silicon nitride nanomembrane. **b.** Exploded view of flow insert layers: L1 = polydimethylsiloxane (PDMS), L2 = pressure-sensitive adhesive (PSA), L3 = polyethylene terephthalate (PET), L4 = PSA. These four layers comprise the flow ports and flow channel. **c.** COMSOL fluid dynamics simulation showing a shear stress map of the flow channel at a flow rate of 500 µl/min. The consistent shear stress over the boxed culture region confirms a uniform flow profile. **d.** Representative 10 x 10 manufactured array of flow inserts, with inset showing a single flow insert unit. Image of fully assembled fluidic hToC, demonstrating phase imaging clarity of endothelial cell monolayer cultured in the flow channel. **e.** Relative cell viability as measured by XTT assay of ECs cultured with insert-conditioned EGM-2 or X-VIVO media. Absorbance normalized to untreated negative control. We observed no differences in cell viability between the treatment conditions as tested by one-way ANOVA, confirming the biocompatibility of the flow insert materials.

Building on our previous approach^[20]^, we adapted our design to have a geometry compatible with commercial manufacturing and characterized flow throughout the device in COMSOL multiphysics software for target shear stress at the culture surface ranging from 1–10 dynes/cm^2^. The shear stress map **(Fig. 1c)** confirmed a uniform flow profile across the culture region and defined flow rates that achieve physiological shear stress levels in the device. Based on this design, prototype flow inserts were constructed in-house with polydimethylsiloxane (PDMS) via standard soft lithography techniques. These fully PDMS-based components were sterilized with ethanol and the surface was activated with UV-ozone to enable adhesion to the membrane chip. After a 24-hour curing period, the devices were fully sealed for fluidic experiments. Proving to be leak-proof and biocompatible, these ‘in-house’ flow inserts validated the basic design concept for advancement to manufacturing at ALine. To confirm biocompatibility and ensure the absence of cytotoxic leachates from the manufactured components, we performed an XTT viability assay in which in-house fabricated or commercially manufactured inserts were incubated in media types used in our tissue chip model (EGM-2, X-VIVO). The conditioned media was then added to human umbilical vein endothelial cells (ECs) in a 96-well plate, and the cells were cultured for 24 hours before measuring cell viability. In both endothelial growth media and serum-free X-VIVO, we observed no differences in cell viability between the manufactured insert-treated group and the control group **(Fig. 1e)**.

The final commercially manufactured flow insert **(Fig. 1a, b)** is composed of a 3 mm thick PDMS body with flow ports connected to a 200 μm tall flow channel. The flow channel comprises two PSA layers that encompass a polyethylene terephthalate layer. The total volume including the ports and channel is 4.23 μl **(Supplementary Table S1)**. Parts are manufactured in a 10 x 10 array with bridging gates between individual inserts that are cut out upon receipt in the lab **(Supplementary Fig. S1).** The presence of a bottom PSA layer enables the desired peel-and-stick functionality with sealing and alignment achieved using the same jigs as for the hToC assembly. Once assembled in the hToC device, the flow insert’s channel encompasses the porous membrane to enable flow across the culture region **(Supplementary Fig. S2)**. In our initial validation, we confirmed the efficiency of assembly (<5 min), leak-proof performance (≥99%), and optical transparency for live cell imaging by phase contrast microscopy **(Fig. 1d)**.

### Demonstration of Shear Stress Dependent EC Alignment in Manufactured Inserts

Shear stress has been extensively shown to induce elongation and remodeling of actin fibers aligned in the flow direction.^[17,20]^ To validate that the manufactured flow inserts reproduce characteristic EC morphological alignment, we first tested EC biocompatibility by statically culturing ECs within the flow channel for 24 hours. Live/dead staining revealed comparable cell viability (97.7 ± 0.6%) to that of the prototype in-house inserts (96.0 ± 1.0%) **(Fig. 2b)**.

**Figure 2.**
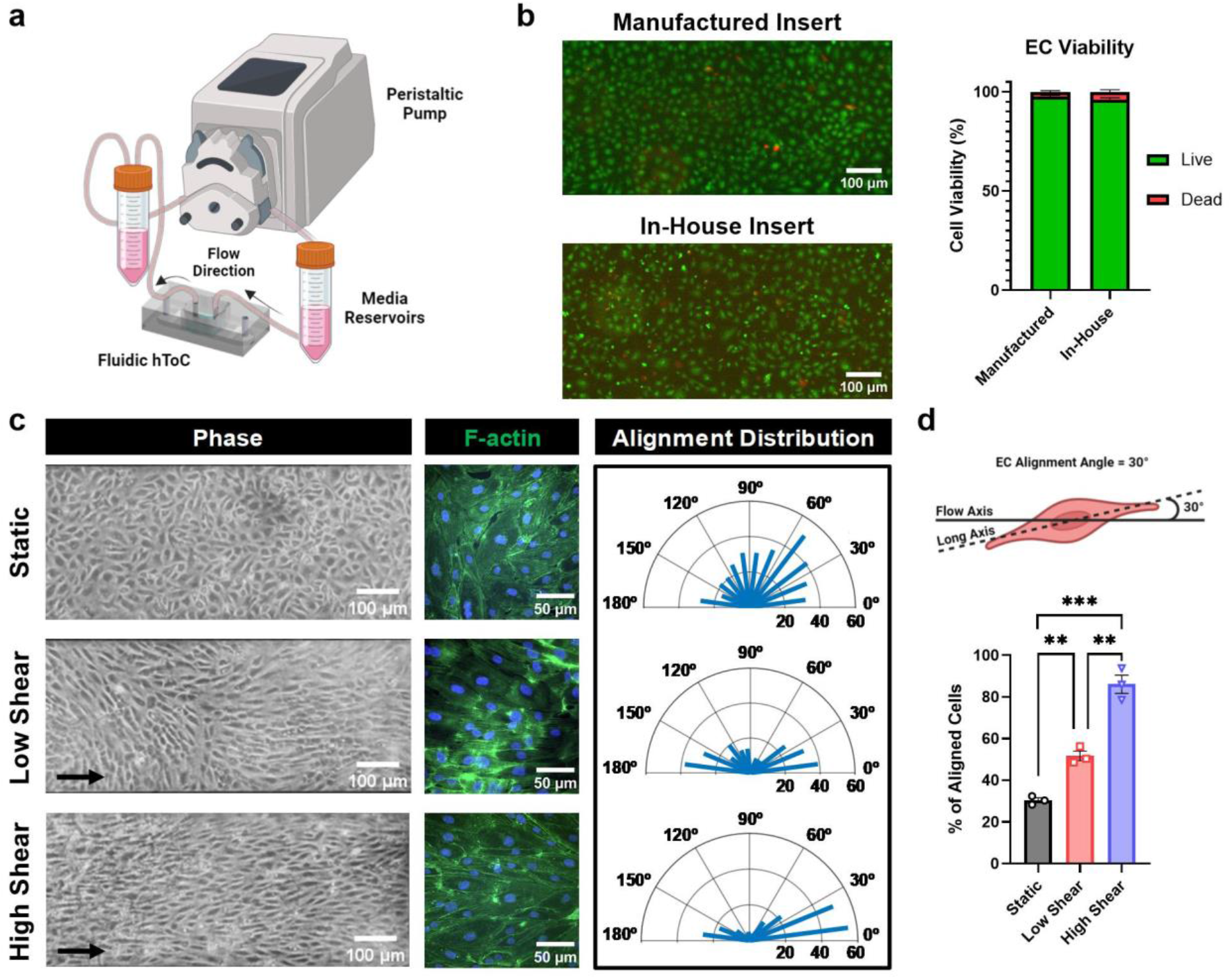
Flow insert is biocompatible with endothelial culture and reproduces physiological cytoskeletal remodeling with fluidic shear stress. **a.** Peristaltic pump flow setup containing media reservoirs that serve as pulse dampeners are connected to the fluidic hToC device. **b.** Live/dead assay results of ECs cultured in either the manufactured flow insert or in-house insert showing comparable viability after 24 hours of static culture in the flow channel (Live = green, Dead = red). **c.** Phase images of ECs cultured in fluidic hToC for 24 hours of either static culture or physiological fluidic shear stress of 1.5 or 10 dynes/cm^2^. Black arrows indicate direction of flow. ECs were stained for F-actin to visualize cytoskeletal remodeling in response to flow. Radar plots quantify cell alignment relative to the flow direction. **d.** Schematic demonstrating the definition of alignment angle, which is calculated as the angle between the flow axis and long axis of the cell. Quantification of percent of aligned cells, where alignment angle <30° or >150° is considered aligned along the flow axis. Increasing shear stress resulted in an increasing number of aligned cells. One-way ANOVA with Tukey’s post-hoc test was used, **p < 0.01, ***p < 0.001.

For flow experiments, we used a peristaltic pump circuit with fluidic capacitors to dampen oscillations and produce non-pulsatile flow^[24]^ **(Fig. 2a)**. ECs were cultured for 24 hours either statically or under fluidic shear stress of 1.5 or 10 dynes/cm^2^.^[14,15]^ We selected a low shear condition of 1.5 dynes/cm^2^ to represent microvascular flow in the postcapillary venule and a high shear condition of 10 dynes/cm^2^ to reflect typical capillary flow. Phase contrast imaging on the optically transparent membranes demonstrated cell elongation, further supported by immunofluorescence staining of F-actin **(Fig. 2c)**. Radar plots illustrating the distribution of aligned cells across the culture region demonstrated a greater proportion of aligned cells in fluidic conditions compared to static cultures. We defined alignment angle as the angle between the long axis of the cell and the flow axis, and cells with an alignment angle <30° or >150° were categorized as aligned. Quantification in Figure 2d revealed a shear-stress dependent increase in the percentage of aligned cells (30.4 ± 2.1% for static, 48.6 ± 4.0% for low shear, 78.7 ± 7.6% for high shear). These results established the capacity of the manufactured inserts to produce physiologically relevant shear stress-dependent cell alignment. Based on these results, we next sought to examine the consequences of fluid flow on cell physiology in the hToC, beginning with the molecular responses of endothelial cells.

### Vascular Flow Improves Barrier Integrity and Influences Surface Adhesion Protein Expression

Cell–cell junctions regulate endothelial barrier permeability to both solutes and circulating immune cells.^[25]^ Junctional integrity depends on the function of adhesive proteins, such as VE-cadherin and PECAM-1, which are influenced by shear stress at the vascular wall.^[26]^ In ECs cultured with 1.5 dynes/cm^2^ shear stress, we observed a decrease in VE-cadherin gap width **(Fig. 3b)** and morphology that was distinct from the continuous appearance in static cultures **(Fig. 3a, inset)**. The irregular junction lines in fluidic cultures are a morphological adaption to shear stress, as reported in other studies^[27,28]^. Additionally, we found that PECAM-1 staining in fluidic conditions was more consistent across the cell monolayer contrasting with the patchy and discontinuous expression observed in static cultures **(Fig. 3a)**. Given the role of PECAM-1 in leukocyte transmigration,^[29]^ its more uniform expression under flow may indicate an adaptive response to shear stress, which is important for improving barrier properties and enabling leukocyte trafficking.

**Figure 3.**
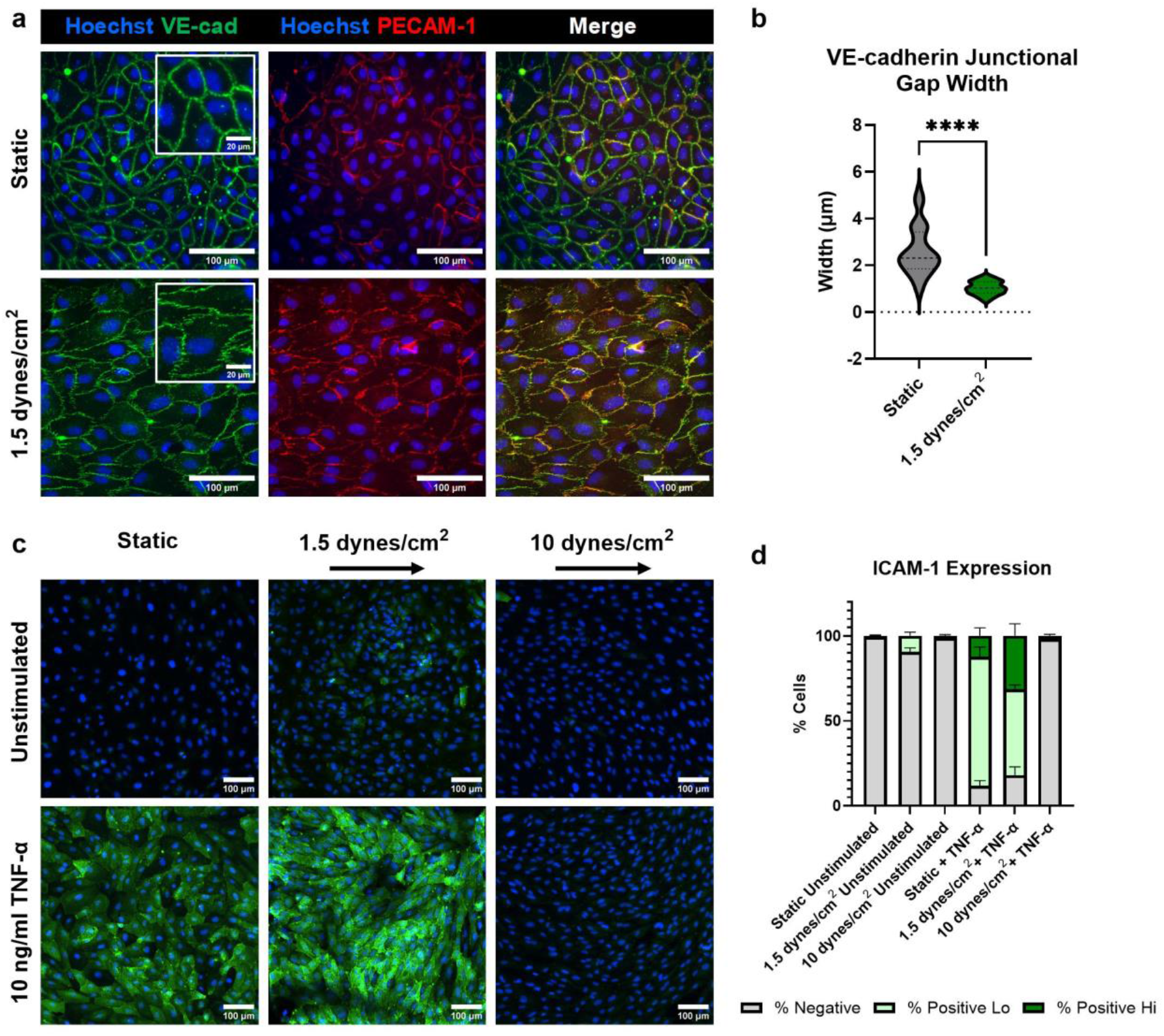
Effect of fluid flow on surface and junctional adhesion molecule expression with or without TNF-α stimulation. **a.** ECs cultured for 72 hours, either statically or with 1.5 dynes/cm^2^ of fluidic shear stress. Junctions labeled with VE-cadherin (green) and PECAM-1 (red). Insets show an enlarged view of the VE-cadherin morphological differences between fluidic and static cultures. **b.** Quantification of VE-cadherin junctional gap width, demonstrating a significant decrease in gap width for fluidic cultures. Unpaired t-test was used to determine statistical significance, ****p < 0.0001. **c.** ECs cultured with basal 10 ng/ml TNF-α stimulation for 24 hours of static culture or fluidic shear stress of 1.5 or 10 dynes/cm^2^. ECs were preconditioned with 1.5 dynes/cm^2^ of shear stress for 24 hours before stimulation. Devices were stained with ICAM-1 (green) and Hoechst (blue). **d.** ICAM-1 expression is reported as the percent of cells with high, low, or no ICAM-1 levels. A fluorescence intensity threshold was set across all images to determine expression levels. Results indicate that a greater percentage of ECs express high levels of ICAM-1 in the 1.5 dynes/cm^2^ condition with TNF-α stimulation compared to static culture (p < 0.001, one-way ANOVA with Tukey’s post-hoc test).

Next, we investigated the impact of preconditioning with flow on the inflammation-driven expression of the surface adhesion molecule ICAM-1, which mediates immune cell adhesion and transmigration.^[30]^ For these experiments, we used TNF-α, a prominent cytokine in the tendon inflammatory milieu that promotes the upregulation of ICAM-1 on HUVECs.^[31,32]^ ICAM-1 intensity levels were measured with immunocytochemistry and compared across six experimental groups: flow (24 hours at low (1.5 dynes/cm^2^) or high (10 dynes/cm^2^) shear stress) or statically cultured with and without TNF-α stimulation. Our results demonstrated that both static and fluidic cultures maintained low baseline ICAM-1 expression in the absence of cytokine stimulation **(Fig. 3c)**. However, TNF-α-stimulated ECs under low shear stress (1.5 dynes/cm^2^) accentuated ICAM-1 expression while high shear stress (10 dynes/cm^2^) attenuated ICAM-1 to levels comparable to unstimulated controls. The high shear stress results align with previous studies^[33]^, in which TNF-α-stimulated HUVECs subjected to 15 dynes/cm^2^ of fluidic shear stress exhibited a significant decrease in ICAM-1 intensity 24 hours of flow. Interestingly, under low shear stress conditions with stimulation, we found that the total percentage of ICAM-1-positive cells (82.0 ± 4.3%) was comparable to that under static-stimulated conditions (88.3 ± 2.7%) **(Fig. 3d)**. However, when we further defined an intensity threshold of high versus low ICAM-1 expression to capture the heterogeneity across the monolayer, a significantly greater (p < 0.001) proportion of ECs displayed high levels of ICAM-1 expression under low shear stress conditions (31.5 ± 6.2% for low shear, 12.3 ± 4.1% for static; **Fig. 3d)**. These findings highlight the complex interplay between flow-induced shear stress and inflammatory signaling, informing our subsequent studies on monocyte transmigration across the inflamed endothelium in the hToC.

### Circulation Enables Modeling of the Monocyte Transmigration Cascade

Monocyte infiltration from the circulation plays a central role in the normal immune response following injury, but overactivation contributes to the development of tendon fibrosis.^[34]^ While our static hToC captured the role of infiltrating monocytes, the absence of physiological flow impacts endothelial-monocyte interactions.^[35,36]^ To address this, we next developed a method to introduce circulating monocytes through the vascular flow channel. Since live imaging requires maintaining flow outside of the controlled environment of the incubator, we optimized a flow setup for live imaging to monitor the circulating monocytes in real time, which included a bead bath for the media reservoirs and an incubation stage to maintain humidity at 95% and temperature at 37°C **(Fig. 4a)**.

**Figure 4.**
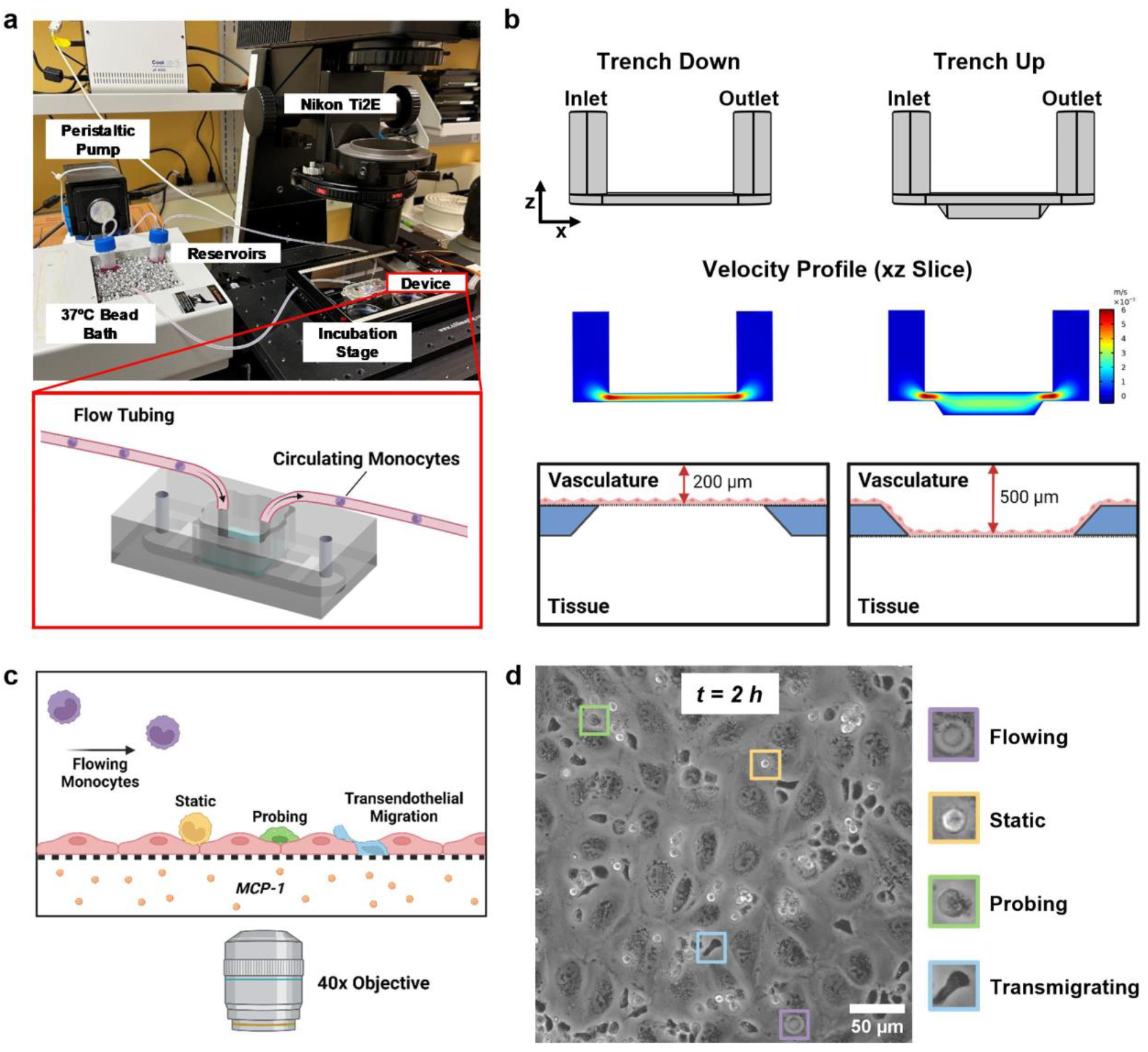
Establishing a live imaging setup to capture circulating monocyte transmigration with a ‘trench up’ membrane configuration. **a.** Live imaging flow setup, which is comprised of a peristaltic pump, media reservoirs in a 37°C bead bath, and an incubation stage that holds the device. This setup sits on a Nikon Ti2E microscope to capture real-time phase videos. Schematic of fluidic hToC with circulating monocytes. **b.** Comparison of the flow channel geometry between ‘trench down’ and ‘trench up’ membrane configurations. Velocity profiles of an xz slice through the center of the channel, demonstrating a decreased flow velocity within the trench. **c.** Schematic illustrating the steps in the monocyte transmigration cascade, which include static adhesion to the endothelium, probing, and transmigration across the barrier. (MCP-1: Monocyte chemoattractant protein-1) **d.** The final frame of a 2-hour live video of monocyte circulation showing the EC monolayer with monocytes at various stages along the transmigration cascade. Color coding corresponds to the monocyte state along the transmigration cascade based on the phase appearance of the cells. Monocytes that appear phase bright are on top of the endothelium, cells that are flattening out and probing appear gray, and cells that have transmigrated under the endothelium appear phase dark.

In previous experiments, the membrane chips are oriented ‘trench down’, which creates a continuous channel across the porous window and surrounding regions **(Fig. 4b)**. With shear stresses ranging between 1.5–4.5 dynes/cm^2^, the postcapillary venule is the primary site of immune cell transmigration, where monocytes exit the circulation and enter the injured tissue.^[15]^ In the initial inflammatory stages of injury, a transient increase in vascular permeability leads to hemoconcentration which slows blood flow locally and enables leukocytes to gain access to the endothelium.^[37]^ To mimic the slowing of blood flow at the postcapillary venule, we reoriented the chip to a ‘trench up’ configuration, which increases the vascular channel height by 2.5 times **(Fig. 4b)**. We performed COMSOL simulations to model the altered shear stress profile **(Supplementary Fig. S3)** and defined a maximum flow rate to avoid off-target leukocyte activation, which occurs above 1.5 dynes/cm^2^.^[38,39]^ A table summarizing ‘trench down’ and ‘trench up’ shear stress values across a range of flow rates (10–1000 µl/min) is provided in **Supplementary Table S2**.

With these characterizations in place, we circulated human monocytes isolated from healthy donors at a flow rate of 50 μl/min which maintained a maximum shear stress throughout the channel below 1.5 dynes/cm^2^. The monocyte concentration in the flow circuit was optimized at 200,000 cells/ml according to representative levels of normal human blood counts.^[40]^ We leveraged real-time phase microscopy, enabled by the glass-like imaging clarity of the membrane and transparency of the flow insert, to validate the monocyte transmigration cascade under flow in response to the tissue-side addition of MCP-1 over 2 hours. The multistep transmigration cascade **(Fig. 4c)**, regulated by shear flow, basal chemokines, and inducible adhesion molecules expressed by the endothelium, was captured in real time.^[32]^ The 2-hour time-lapse video **(Supplementary Video 1)** demonstrated key aspects of physiological monocyte transmigration, including rolling, arrest to the endothelial surface, and crossing of the endothelial barrier. Using phase imaging, we classified cells at different stages of the transmigration process^[41,42]^: phase-bright cells are on top of the endothelium, gray cells are flattening out and probing, and phase-dark cells have transmigrated under the endothelium **(Fig. 4d)**. This analysis confirmed that the circulating system enabled by the manufactured flow insert successfully mimics the multistep interactions of immune cells with the vascular wall.

### Circulation Facilitates Robust Monocyte Infiltration Over a 24-Hour Timescale

Since the complete hToC disease model is cultured for 24 hours, we extended the timescale of monocyte circulation by live-labeling the monocytes and quantifying transmigration into the bottom channel with or without MCP-1. Confocal image reconstructions show an xz view of the devices **(Fig. 5a)**, where monocytes adhering to the endothelium are highlighted in white and transmigrated monocytes are highlighted in orange. To quantify monocytes across the 3D volume, we utilized a classification tool in Imaris **(Supplementary Fig. S5)**. We observed a striking increase in the number of transmigrated monocytes in the circulating condition with MCP-1, while we found a non-significant response to MCP-1 in static culture conditions **(Fig. 5b)**. Additionally, we observed a significant increase (p = 0.0248) in the number of adhered monocytes in the circulating +MCP-1 condition, with non-significant differences in static culture. These results are consistent with reports that pre-sheared endothelial monolayers and leukocyte rolling under flow induce a migratory leukocyte phenotype.^[43]^ The addition of flow may also create a steep chemotactic gradient between the top and bottom channels, enhancing the monocyte transmigration response and underscoring the importance of circulation in this context. As part of our studies, we also compared monocyte transmigration across endothelium seeded on dual-scale membranes with micropore sizes of 3 μm or 5 μm. Previously, we used 3 μm and 5 μm membranes for neutrophil^[31,41]^ and monocyte^[10]^ transmigration studies, respectively. Here, we found no significant differences in monocyte transmigration between these two micropore sizes **(Supplementary Fig. S5)**, enabling the use of 3 μm membranes in future monocyte studies.

**Figure 5.**
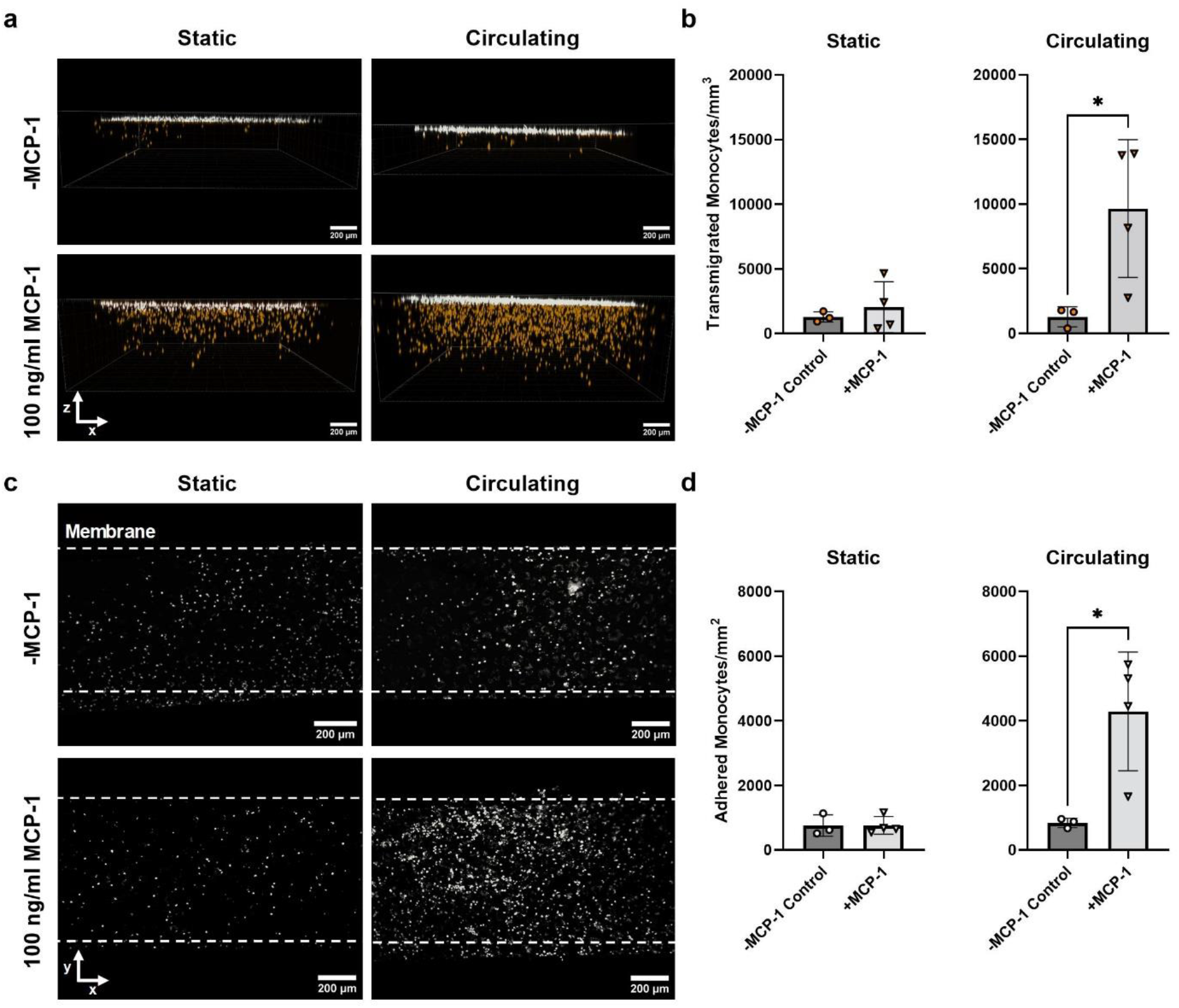
Monocyte circulation facilitates increased transmigration and adhesion compared to static conditions. **a.** 3D volume reconstruction of confocal image stacks showing adhered (white) and transmigrated (orange) monocytes after 24 hours of either static culture or circulation (50 μl/min) in fluidic hToC devices with endothelial barriers. 100 ng/ml MCP-1 was added to the bottom channel and compared to a (-) MCP-1 control condition. **b.** Quantification of transmigrated monocytes, normalized over the total image volume. Results demonstrate increased levels of monocyte transmigration in response to MCP-1 in the circulating condition, compared to non-significant changes in static culture. **c.** xy slice at the membrane displaying monocytes adhered to the endothelium **d.** Quantification of adhered monocytes, normalized over the total image area. Circulating flow resulted in a significant increase (p = 0.0248) in monocyte adhesion. One-way ANOVA with Tukey’s post-hoc test was used, *p < 0.05.

### Monocyte Transmigration and Endothelial Activation in Response to Cytokine Stimulation Under Flow

To further dissect the circulating monocyte response to cytokines in the fibrotic secretome, we quantified monocyte transmigration across cytokine-stimulated ECs. ECs were preconditioned at 1.5 dynes/cm^2^ for 24 hours, followed by the addition of TNF-α or TGF-β1 to the bottom channel with flow continued for another 24 hours. Monocytes were then introduced at a flow rate of 50 µl/min and were recorded over 2 hours using phase contrast microscopy to visually assess transmigration. Under control conditions, monocytes appeared phase bright with a rounded morphology **(Fig. 6a)**. In contrast, 63.9±11.4% of monocytes appeared phase dark with an elongated morphology in response to the TNF-α-stimulated endothelium **(Fig. 6a, b)**. Interestingly, TGF-β1 stimulation did not induce a pro-migratory response and we observed a greater proportion of monocytes that remained phase bright, with only 24.1±8.2% transmigration. We additionally compared percent transmigration in response to MCP-1, a potent monocyte chemoattractant that is highly upregulated in the hToC milieu.^[10]^ As expected, 55.4±5.0% of the monocytes transmigrated, which was similar to what was observed in the TNF-α group. To investigate whether MCP-1 and TGF-β1 induce a combinatory transmigration response, we added MCP-1 to TGF-β1-stimulated ECs before monocyte addition. TGF-β1+MCP-1 resulted in transmigration levels similar to those of MCP-1 alone **(Fig. 6b)**, indicating that MCP-1 is primarily responsible for driving transmigration.

**Figure 6.**
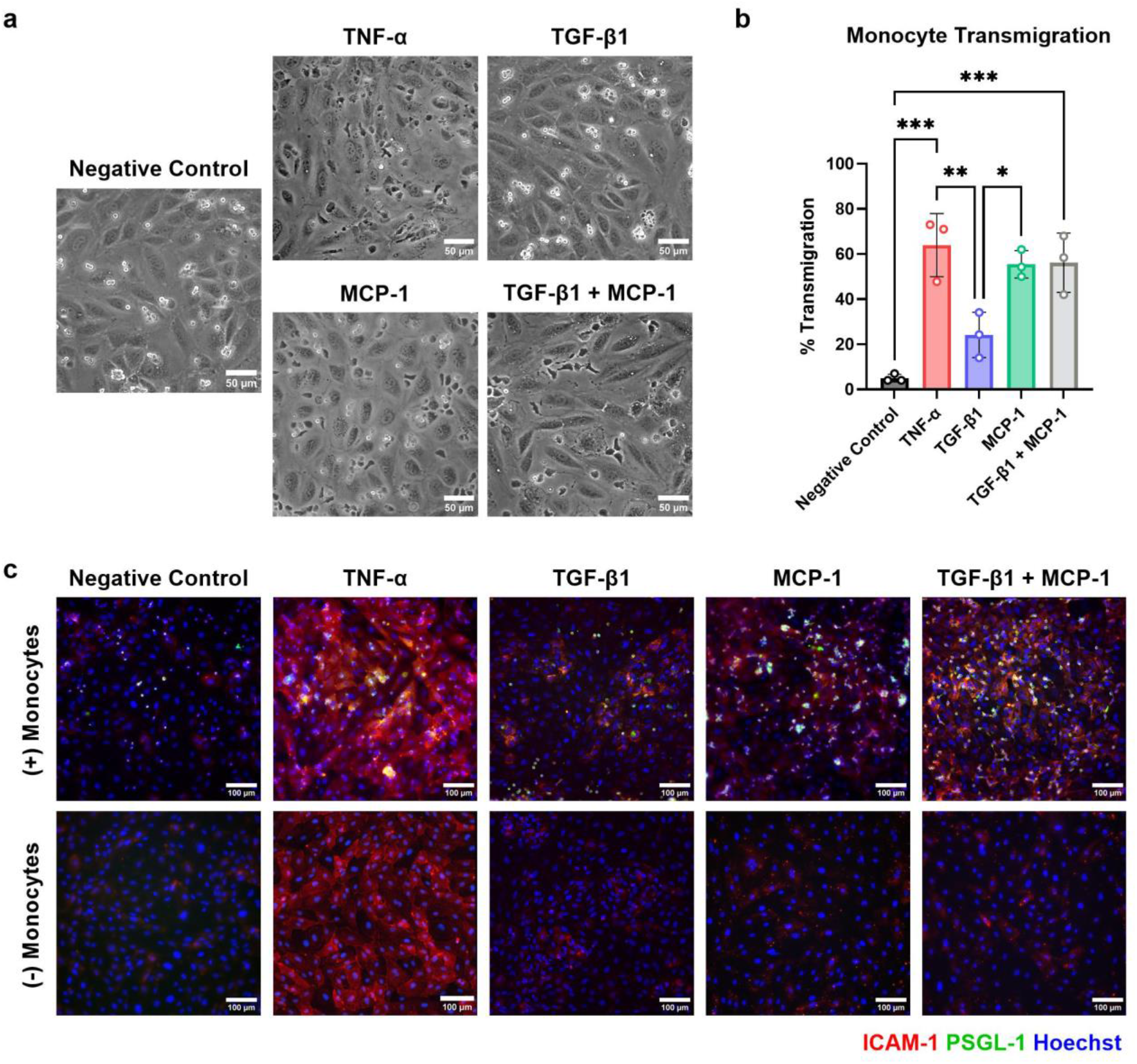
Monocyte transmigration and endothelial inflammatory response to key cytokines in the fibrotic secretome. **a.** Monocytes circulated in EC-seeded fluidic hToC devices were observed in phase contrast time-lapse recordings over 2 hours. Representative images of the final frame of the 2-hour videos are shown. **b.** Percent transmigration quantification of N = 3 experimental replicates per condition. Results demonstrated significantly elevated monocyte transmigration levels in response to TNF-α stimulated ECs compared to the negative control (p < 0.001) and TGF-β1 stimulation condition (p = 0.005). Similarly, MCP-1 and TGF-β1+MCP-1 induced significantly increased percent transmigration compared to the negative control (p < 0.001) and TGF-β1 stimulation condition (p = 0.020). One-way ANOVA with Tukey’s post-hoc test was used, *p < 0.05, **p < 0.01, ***p < 0.001. **c.** Following the 2-hour transmigration experiment, devices were stained for ICAM-1 (red), PSGL-1 (green), and Hoechst (blue). Representative images of devices with and without monocytes in each stimulation condition are shown. As expected, we observed a substantial increase in ICAM-1 upon TNF-α stimulation regardless of the presence of circulating monocytes. The MCP-1, TGF-β1, and TGF-β1+MCP-1 conditions, on the other hand, did not induce ICAM-1 upregulation. However, upon monocyte circulation, we observed an increase in ICAM-1 expression in these three conditions.

We next assessed the levels of endothelial activation under the tested stimulation conditions (TNF-α, TGF-β1, MCP-1, TGF-β1+MCP-1), both with and without the addition of monocytes, by performing immunocytochemical staining for ICAM-1 and PSGL-1. PSGL-1 is expressed by monocytes and facilitates rolling and attachment to the endothelium via EC selectins.^[44]^ Compared to the control, TNF-α stimulation resulted in a substantial increase in ICAM-1 expression regardless of monocyte introduction **(Fig. 6c)**. Consistent with the results of the transmigration quantification, TGF-β1 induced minimal ICAM-1 expression. Although MCP-1 alone did not upregulate ICAM-1, it led to increased monocyte adhesion (as shown by PSGL-1 staining) and high transmigration rates, which together resulted in EC activation. Similarly, we observed a marked increase in ICAM-1 in devices with circulating monocytes in the TGF-β1+MCP-1 condition. Overall, these results indicate that strong EC activation contributes to monocyte transmigration and that monocyte transmigration can further lead to EC activation.

### Tissue Macrophages Increase Monocyte Infiltration in the Tendon-Vascular Disease Model

Once we established a basic understanding of flow on ECs and monocyte transmigration, we sought to study its impact on the complete hToC model. This culture system includes a top vascular compartment with ECs and circulating monocytes, as well as a bottom tissue compartment composed of tendon fibroblasts (tenocytes) and tissue macrophages (tMφ) encapsulated in a hydrogel. To incorporate flow, we established a culture timeline to ensure preconditioning of the endothelial monolayer and the introduction of circulating monocytes under flow **(Fig. 7a)**. First, to establish a tissue-side culture, tenocytes and blood-derived monocytes were added to a type I collagen hydrogel at D_-6_ and cultured in serum-free X-VIVO media spiked with M-CSF to differentiate the monocytes into a naïve macrophage state. At D_-2_ of the timeline, ECs were seeded through the flow insert, and the top component was mounted on a media reservoir according to previously established methods.^[10]^ After 24 hours of static culture to establish the EC monolayer, the tissue and vascular components were combined, and 1.5 dynes/cm^2^ shear stress was applied to the vascular channel. At D_0_, monocytes were added to the flow circuit, and the flow rate was reduced to 50 μl/min to avoid off-target activation.

**Figure 7.**
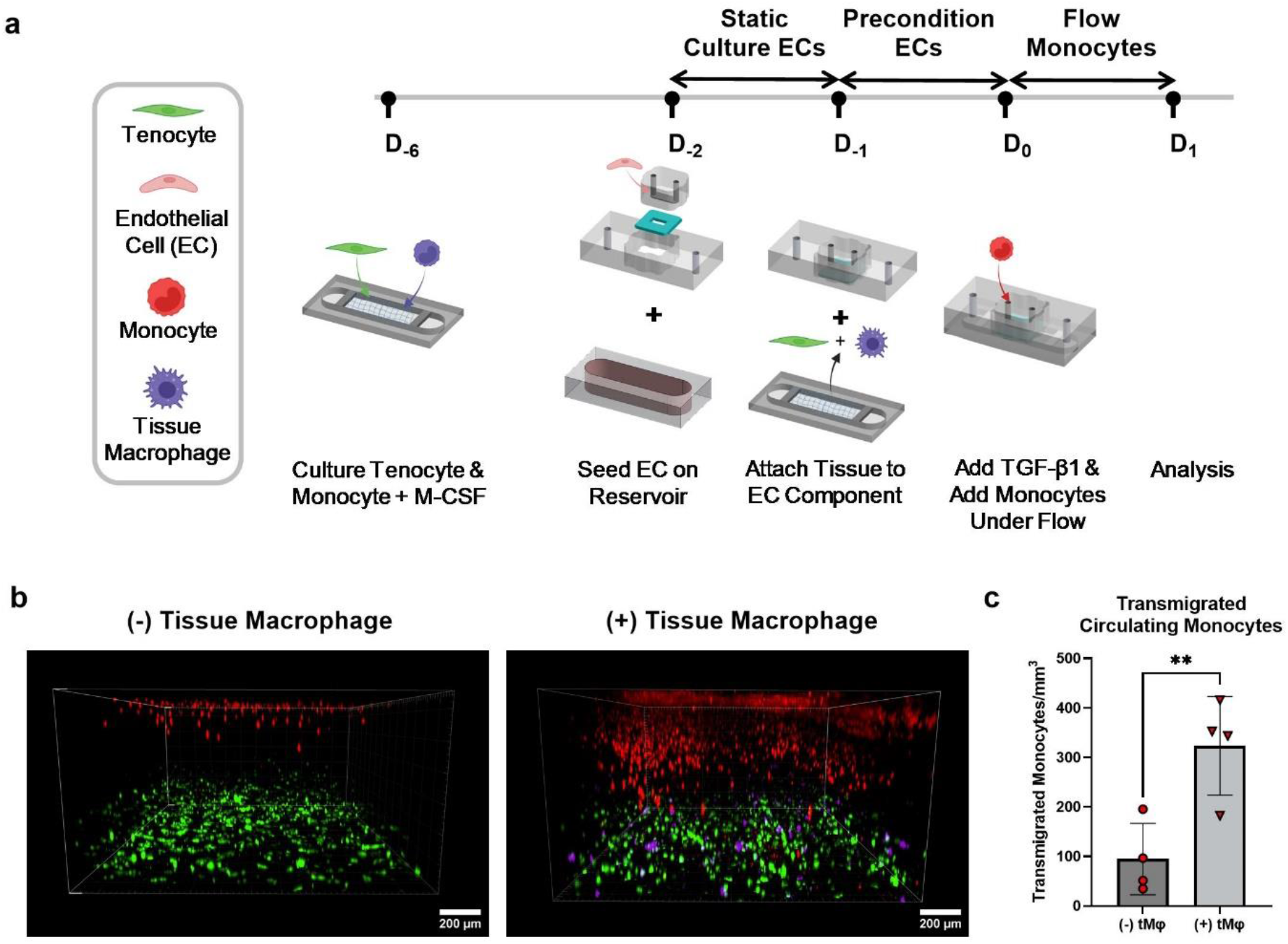
Integration of vascular flow in the complete hToC culture. **a.** Culture timeline for complete fluidic hToC model, including all four cell types (tenocytes, ECs, monocytes, and tissue macrophages). Tenocytes and monocytes were encapsulated in a collagen hydrogel and cultured in the presence of macrophage colony-stimulating factor (M-CSF) for 6 days. ECs were statically cultured in the vascular flow channel for 24 hours and then preconditioned at 1.5 dynes/cm^2^ for 24 hours. At this point, monocytes were added to the circulation for an additional 24 hours. **b.** Confocal images of hToC cultures with or without tissue macrophages (tMφ) in the tissue hydrogel after 24 hours of monocyte circulation through the vascular channel (D_1_). All cells other than the EC monolayer were live-labeled with membrane dyes. Circulating monocytes are red, tenocytes are green, and tissue macrophages are purple. The presence of tissue macrophages drives circulating monocyte transmigration into the bottom channel. **c.** Quantification of transmigrated circulating monocytes, showing a significant increase (p = 0.0098) in infiltration with the presence of tissue macrophages. One-way ANOVA with Tukey’s post-hoc test was used, **p < 0.01.

To validate this timeline, we live-labeled tenocytes, macrophages, and circulating monocytes to visualize the culture at 24 hours. Confocal volume reconstruction of the 3D culture environment with or without tMφ in the bottom channel hydrogel showed circulating monocytes, tenocytes, and tMφ labeled in red, green, and purple, respectively **(Fig. 7b)**. In the absence of tMφ, monocyte transmigration was limited, with <100 transmigrated monocytes/mm^3^. In contrast, the presence of tMφ significantly increased (p = 0.0098) monocyte transmigration, with >300 transmigrated monocytes/mm^3^ **(Fig. 7c)**. This difference suggests that tissue macrophages play a critical role in promoting monocyte transmigration, likely through secreting factors such as MCP-1 and TNF-α to promote the transmigration process.

## Discussion

The influx of vasculature and infiltration of immune cells following tendon injury shifts the repair process toward fibrosis.^[7]^ To understand vascular crosstalk within the tendon microenvironment, our group developed a barrier tissue system to model the tendon-vascular interface. However, the model’s static vascular component does not include the crucial physiologic factor of flow, which impacts endothelial barrier function and downstream inflammatory responses. In this work, we introduced and validated a fluidic human tendon-on-a-chip (hToC) to investigate endothelial-monocyte interactions under physiological shear stress, replicating vascular inflammatory fibrosis. We developed a novel peel-and-stick flow component for easy assembly and commercial scalability. Our initial validation studies confirmed that the flow insert accurately mimics physiological shear stress, supports endothelial cell viability, and induces EC elongation due to cytoskeletal remodeling^[17]^. Using this validated system, we found that shear stress improved endothelial barrier integrity, with VE-cadherin and PECAM-1 reorganization. ICAM-1 expression on TNF-α stimulated ECs increased under low shear (1.5 dynes/cm²) but was reduced at higher shear (10 dynes/cm²). Monocyte transmigration was significantly enhanced by TNF-α and MCP-1 under flow, and tissue macrophages further amplified monocyte infiltration into the tissue with circulating flow. These results demonstrate the model’s utility in studying vascular inflammation and immune cell transendothelial migration dynamics in tendon fibrosis.

While we confirmed that flow improved endothelial barrier properties, we observed an intriguing relationship between shear stress magnitude, inflammatory challenge, and ICAM-1 expression. With low shear stress and TNF-α stimulation, the number of ECs with high levels of ICAM-1 expression was markedly greater than that of static stimulated ECs. These results align with vascular physiology where a reduced shear environment in postcapillary venules is the site of ICAM-1 mediated recruitment of leukocytes.^[45]^ In contrast, ICAM-1 expression was abrogated at high shear levels under inflammatory conditions, returning levels to baseline. We also investigated whether a lack of EC preconditioning with shear was important for preventing ICAM-1 upregulation under inflammatory conditions at high shear (10 dynes/cm^2^). Even without preconditioning, we found that high shear results in baseline levels of ICAM-1 expression as seen in static cultures **(Supplementary Fig. S4)**. Chiu *et al.* reported similar anti-inflammatory effects on the EC response to inflammatory stimuli with and without preconditioning at 20 dynes/cm^2^.^[46]^

Infiltrating immune cells from the vasculature, specifically monocytes, play a significant role in perpetuating the inflammatory response following tendon injury.^[47]^ As the vasculature transitions from the capillary bed to the venule, there is a doubling in vessel diameter as well as a slowing of blood flow to enable immune cell engagement with the endothelium during inflammation. To simulate this geometric expansion from the capillary to the postcapillary venule, we leveraged a unique feature of the ultrathin nanomembranes. Inverting the chip compared to our prior work^[10,18,24,38]^ resulted in an expansion of the flow channel height from 200 μm to 500 μm directly above the membrane, enabling immune cell engagement and migration across the vascular barrier. While EC and junction remodeling in response to fluidic shear conditioning suggests improved barrier integrity, we also observed robust increases in monocyte transmigration under these same conditions. This apparent paradox can be explained by the maintenance of a sharp chemotactic gradient in fluidic conditions^[48]^ as well as increased MCP-1 presentation by the EC glycocalyx, which is upregulated in response to flow^[49]^. In the postinjury response *in vivo*, chemokines including MCP-1 are secreted by ECs, fibroblasts, and macrophages and play important roles in selectively recruiting monocytes from the bloodstream. Tissue-secreted MCP-1 is presented on the apical EC surface via the glycocalyx and is immobilized by ECM proteins on the basal side to preserve the chemotactic gradient.^[50,51]^

Interestingly, we demonstrated that MCP-1 alone does not lead to EC activation, despite it producing a high degree of monocyte transmigration. This was most notable in the MCP-1 and TGF-β1 + MCP-1 conditions, where the basal introduction of MCP-1 did not upregulate ICAM-1 expression on the endothelium. With the addition of monocytes to the MCP-1 and TGF-β1 + MCP-1 conditions, we observed monocytes expressing PSGL-1 as well as ICAM-1 upregulation. These observations can be explained by the role of PSGL-1 in triggering inflammation and increased ICAM-1 expression^[52]^. To the best of our knowledge, no studies have directly investigated the effect of MCP-1 on ICAM-1 expression in human ECs, although the functions of MCP-1 extend beyond chemoattraction.^[53]^ It has been reported that IL-1 and IL-6 are upregulated by human monocytes upon MCP-1 stimulation^[54,55]^, both of which have been implicated in the upregulation of ICAM-1 on ECs.^[56,57]^ By further analyzing the secretome, we can elucidate the cytokines involved in the ICAM-1 upregulation we observed upon monocyte circulation. Maus *et al.* reported that MCP-1 increases monocyte adhesion to TNF-α-stimulated ECs under flow.^[58]^ In contrast to these findings, we did not observe a co-stimulatory effect of TGF-β1 and MCP-1 in the co-culture conditions explored here. These results likely point to TGF-β1 as a modulator of tissue-sided responses, a question that we probed with the addition of tendon cell types in the bottom channel of the fluidic hToC.

We ultimately incorporated circulating monocytes in the complex cellular microenvironment of the complete hToC model. We found that the presence of tissue macrophages drives a more robust circulating monocyte response into the tissue compared to devices without tissue macrophages. These results are consistent with the documented resident macrophage response in the injured tendon, in which macrophages release pro-inflammatory cytokines, including TNF-α, IL-1β, and IL-6.^[59]^ The formation of a chemokine gradient leads to the recruitment and infiltration of monocytes into the injury site. With these advancements in place, we can test hypotheses about the specific contributions of each cell type to the inflammatory microenvironment in tendon fibrosis. Future work will extend vascular flow studies in the complete hToC model to investigate myofibroblast differentiation, matrix deposition, secretome profiles, and macrophage polarization in the tissue. The inclusion of circulating flow in the hToC will also enable future studies to model systemically administered therapeutics and evaluate drugs targeted against immune cell infiltration and myofibroblast activation.

## Methods

### Nanomembranes

Silicon nitride nanomembranes manufactured by SiMPore, Inc. (Henrietta, NY) were used as substrates to culture endothelial cells in the hToC device. Both nanoporous and dual-scale membranes were used in this study. Nanoporous membranes are ∼100 nm thick, contain ∼60 nm diameter pores, and have a porosity of ∼15%. Dual-scale membranes have the same thickness and nanoporosity but additionally contain 5 µm micropores that add an additional porosity of ∼2%.

### Fluidic hToC Components

The top component, bottom component, and flow insert were manufactured at ALine Inc. (Signal Hill, CA) via laser cutting, lamination, and molding processes compatible with mass production (hundreds to tens of thousands). Top and bottom components were shipped as single units, while the flow inserts were shipped as 5×5 arrays. Individual flow inserts were removed from the array by cutting connecting gates with a razor blade. The flow insert and top component contain access ports to the top flow channel and bottom channel, respectively. All ports can be accessed with P20/P200 pipette tips (VWR, 76322-516). Assembly of top and bottom components was performed in a sterile biosafety cabinet as previously described.^[10]^ Briefly, the protective masking layers that maintain component sterility during shipment were removed. The nanomembrane chip was placed on a custom assembly jig in an inverted or non-inverted orientation using notched tweezers. The top component was then placed over the nanomembrane and pressed firmly to form a tight seal. The bottom component was placed in a separate jig to bond it to the top component. Flow inserts were sprayed with 70% ethanol before being placed in the biosafety cabinet channel side up. The assembled top and bottom components along with the flow inserts were sterilized by 20 minutes of UV exposure in the biosafety cabinet. After UV sterilization, the inserts were rinsed 3 times with sterile 1X PBS, and any excess liquid was removed with a pipette. Using straight-tipped tweezers, the protective lining was removed to expose the pressure-sensitive adhesive on the bottom of the insert. The insert was aligned in the open well of the top component and pressure was applied with the flat side of the tweezers around the perimeter of the insert to activate the adhesive (see supplementary Figure 1). The device is now fully sealed and ready for cell seeding and flow experiments.

A custom peristaltic pump circuit was assembled as previously described.^[20]^ The components included 0.02” ID silicone tubing (VWR, MFLX95802), 21 gauge NT dispensing tips (Jensen Global, JG21-1.0HPX, JG21-1.5HPX-90), and 5 ml vials (BKMAMLAB). A peristaltic pump was used for circulating flow (Langer Instruments, USA).

### COMSOL Multiphysics Simulation

A fluid dynamics simulation was performed in COMSOL Multiphysics to model the fluid profile in the flow insert channel. Laminar flow physics was applied to the channel geometry, and a no-slip boundary condition was selected at the walls. The inlet flow rate was set to a constant value and was simulated across a range from 10–1000 µl/min. A pressure P = 0 was applied to the outlet. A physics-controlled mesh was generated and a stationary parametric sweep study was executed to generate shear stress and velocity profiles.

### Cell Culture Protocols

Cryopreserved pooled human umbilical vein endothelial cells (HUVECs) were purchased from Vec Technologies Inc. and expanded in T-25 flasks (Corning Inc., 430639). HUVECs were cultured in EGM-2 Endothelial Cell Growth Medium (Lonza, CC-3162) and used between passages 2 and 6. Cells were maintained under standard culture conditions (37°C, 5% CO_2_, 95% humidity). For device seeding, the backside of the nanomembrane was first coated with collagen (TeleCol-3, Advanced Biomatrix, 5026) and crosslinked in the incubator for 15 minutes. The flow channel and culture surface of the nanomembrane were then coated with 5 µg/cm^2^ of human fibronectin (Bio-Techne, 1918-FN-02M) for 1 hour at room temperature to facilitate cell attachment. HUVECs were then seeded at 40,000 cells/cm^2^ by infusing a cell suspension at 2×10^6^ cells/ml and cultured statically for 24 hours before starting fluid flow or adding inflammatory stimuli.

For the HUVEC shear conditioning studies, the peristaltic pump was set to a flow rate of 50 or 540 µl/min for low (1.5 dynes/cm^2^) and high (10 dynes/cm^2^) shear stress conditions, respectively. Devices were assembled with a trench-down membrane configuration for monocyte studies, and the flow rate was maintained at 50 µl/min (see supplementary Table 1 for shear stress comparisons). For ICAM-1 studies, HUVECs were basally stimulated with 10 ng/ml TNF-α (R&D Systems, 210-TA) for 18 hours. For live monocyte transmigration studies, HUVECs were basally stimulated with either 10 ng/ml TNF-α or 10 ng/ml TGF-β1 (R&D Systems, 240-B-002) for 18 hours prior to monocyte addition. 100 ng/ml of MCP-1 (R&D Systems, 279-MC) was infused into the bottom channel of devices immediately before monocyte addition.

### Peripheral Blood Mononuclear Cell (PBMC) Isolation and CD14+ Monocyte Sorting

Monocytes were isolated from blood drawn from healthy donors with informed consent before each experiment according to an approved protocol by the University of Rochester Institutional Review Board (STUDY0004777). First, the PBMC layer was isolated from whole blood samples via density separation via 1-Step Polymorphs (Accurate Chemical & Scientific Co., Westbury, NY). The PBMCs were washed twice by centrifuging at 350 × g for 7 minutes at 15°C with sterile-filtered wash buffer consisting of HBSS (Gibco, 14175095), 10 mM HEPES (Gibco, 15630080), and 5 mg/ml BSA (Cell Signaling, 9998S). Red blood cells were then lysed by the addition of 1/6X PBS for 1 minute, followed by 4X PBS and a 7-minute spin at 350 × g. An additional wash step as above was performed before the pellet was suspended in isolation buffer consisting of 1X DPBS (Gibco, 14190250), 2 mM EDTA (Gibco, 15575020), and 1 mg/ml BSA. The cell suspension was filtered through a 70 µm cell strainer, spun down at 300 × g for 3 minutes, and suspended in 80 µl of isolation buffer. To sort monocytes, CD14 microbeads (Milltenyi Biotec, 130-050-201) were added to the cell suspension and incubated at 4°C for 15 minutes. A QuadroMACS™ separator was assembled with an LS column (Milltenyi Biotec, 130-042-401). After centrifuging the cells and microbead suspension again at 300 × g for 3 minutes, the cells were suspended in 500 µl of isolation buffer and added to the LS column. CD14+ monocytes were then flushed out of the column and used in experiments within 3 hours of blood collection.

### Fluidic hToC Culture

The complete hToC quad culture was prepared as previously described^[10]^ with modifications for fluid flow (see Fig. 7A). Cells were labeled with DiI (circulating monocytes), DiO (tenocytes), and DiD (tissue macrophages) from the Vybrant Multicolor Cell-Labeling Kit (Invitrogen, V22889). First, tenocytes isolated from tendon tissue under an approved Institutional Review Board protocol (STUDY00004840) were suspended in a collagen hydrogel at 500,000 cells/ml. Freshly isolated monocytes were added to the hydrogel at a 1:7 ratio of monocytes to tenocytes. 100 µl of the hydrogel cell suspension was added to the bottom channel with 200 µl of XVIVO-10 media (Lonza, 04-380Q) supplemented with 20 ng/ml M-CSF (PeproTech, 300-25). Bottom components were cultured for 6 days (D_-6_ - D_0_), and M-CSF media was half-replaced each day. At D_-2_ the top component was seeded with HUVECs as described above. Briefly, the backside of the membrane was coated with collagen, allowed to crosslink an incubator for 20 minutes, and then placed on a reservoir filled with EGM-2 media. Fibronectin was added to the flow channel at 5 µg/cm^2^ and maintained at room temperature for 1 hour for coating. HUVECs were seeded in the flow channel at 40,000 cells/cm^2^ and cultured statically for 24 hours. At D_-1_, both the top and bottom components were removed from the respective reservoirs and combined to form a complete device. Devices were then connected to the flow, and HUVECs were shear-primed at 1.5 dynes/cm^2^. At D_0_ freshly isolated monocytes were added to the flow circuit at 200,000 cells/ml and 10 ng/ml TGF-β1-supplemented XVIVO-10 was added to the bottom channel. The culture was maintained for 24 hours before confocal imaging.

### Cell Viability Assays

To compare cell viability between in-house produced PDMS flow inserts and manufactured flow inserts, a LIVE/DEAD™ Viability/Cytotoxicity Kit (Invitrogen, L3224) was used according to the manufacturer’s protocol. HUVECs were cultured in both inserts for 24 hours of static culture, and percent viability was quantified in FIJI/ImageJ. To confirm that the manufactured inserts do not release cytotoxic leachates, an XTT (2,3-bis-(2-methoxy-4-nitro-5-sulfophenyl)-2H-tetrazolium-5-carboxanilide) benzene sulfonic acid hydrate) Cell Viability Assay (Biotium, 30007) was performed. Both insert types were placed in either EGM-2 or XVIVO-10 media and incubated for 24 hours. The conditioned media was then added to HUVECs cultured in a 96-well plate. After 24 hours of culture, the XTT assay was performed according to the manufacturer’s instructions. The absorbance readings of sample wells were collected at 450 nm and were subtracted by background absorbance at 630 nm.

### Phase Contrast Microscopy Studies

To acquire phase contrast time-lapse videos, a Nikon Ti2E inverted microscope (Nikon Corporation, Tokyo, Japan) with a sCMOS camera (Zyla 4.2, Andor, UK) and a 40x objective was used. To maintain conditions at 37°C devices were placed in an incubation stage (Okolab, Italy), and media reservoirs were placed in a bead bath. A moistened Kim wipe was placed in the incubation stage for humidity. The fluidic imaging setup is shown in Figure 4A. The monocyte suspension was added to the inlet reservoir of the flow circuit at 200,000 cells/ml. Devices seeded with HUVECs were placed in the incubation stage and connected to the flow circuit. Flow was then started at a rate of 50 µl/min. Phase contrast images were taken every 8 seconds for 1 hour and processed in FIJI/ImageJ (NIH, USA).

### Fluorescence Microscopy Studies

To visualize monocyte transmigration over 24 hours, monocytes were live-labeled with DiI. Monocytes were added at a concentration of 200,000 cells/ml to the flow circuit and the flow rate was set to 50 µl/min. For static devices, a rolling infusion of monocytes at 4000 cells/μl was added, resulting in approximately 5000 cells in the channel. MCP-1 (100 ng/ml) was added to the bottom channels of HUVEC-seeded devices. After 24 hours of monocyte circulation, confocal z-stack images were acquired with an Andor spinning disk confocal microscope (Andor Technology, United Kingdom) on Fusion software with a 10x objective and a 1.33 µm step size. Images were processed in Imaris (Oxford Instruments, United Kingdom).

### Immunofluorescence Staining

For HUVEC staining, flow channels were first washed with PBS. ICAM-1 antibody (BioLegend, 353102) was diluted 1:100 in EGM-2 media and incubated in the flow channel for 15 minutes at 37°C. After washing with PBS, the channels were fixed with 4% paraformaldehyde (Invitrogen, I28800) for 10 minutes. Top and bottom channels were washed three times with PBS, and the flow channel was blocked with 5% goat serum (Invitrogen, containing 0.1% Triton X-100 for 30 minutes at room temperature. The secondary antibody was then diluted 1:200 in blocking solution and added to the flow channel for 1 hour at room temperature (see supplementary Table 2 for secondary antibodies). Nuclei were then stained with Hoechst 33342 (Invitrogen, H3570) diluted 1:10,000 in PBS for 5 minutes. For VE-cadherin (R&D Systems, MAB8381) and PECAM-1 (Invitrogen, PA5-32321), antibodies were diluted (1:100) and added after the fixation step. To stain for F-actin, HUVECs were permeabilized with 0.1% Triton X-100 in PBS after fixation. Phalloidin conjugated antibody (Abcam, ab176753) was diluted 1:1000 in 1% BSA and incubated at room temperature for 20 minutes. Images were acquired with an Andor spinning disk confocal microscope on Fusion software. Images were processed in Image and the same brightness and contrast settings were applied to images of all experimental conditions for each channel.

### Image Analysis

For VE-cadherin junctional gap width quantification, 30 measurements were taken in each field of view across three separate devices per condition. For ICAM-1 quantification, an intensity threshold was defined based on the distribution of ICAM-1 signal to determine a high and low ICAM-1 expression cutoff. Cells were segmented in ImageJ to calculate the mean pixel intensity per cell and the percent of cells within each ICAM-1 expression threshold was plotted out of 100%. To quantify cell alignment, cells were stained with a LIVE/DEAD™ kit as above and fitted with ellipses using the Analyze Particles feature of ImageJ. Angles relative to the flow axis per cell were graphed as radar plots in MATLAB using the CircHist plugin.^[60]^ Monocyte transmigration from time-lapse phase videos was quantified manually in ImageJ by counting phase dark and phase bright monocytes in the final frame. To quantify fluorescently labeled monocytes in confocal images, the Spots tool in Imaris was used to identify cells based on a specified diameter. Cells below the membrane were considered to be transmigrated.

### Statistical Analysis

All experiments were performed in triplicate and statistically analyzed with GraphPad Prism software (GraphPad, La Jolla, CA). An unpaired t-test was used to compare differences between two groups and an ordinary one-way ANOVA with Tukey’s post-hoc test was used for comparison of multiple groups, with p < 0.05 considered statistically significant. All quantifiable data are reported as the mean ± standard deviation.

## Data Availability

All data related to the current study are available from the corresponding author upon reasonable request.

## Acknowledgments

I.L., A.S., B.L.M., H.A.A., and J.L.M were supported by NIH UH3TR003281. I.L. was also supported by the National Science Foundation Graduate Research Fellowship. Schematics were created with BioRender.com.

## Author Contributions

Conceptualization: I.L., S.B., H.A.A., J.L.M. Experiments and Data Acquisition: I.L. Data Analysis and Interpretation: I.L., J.L.M., H.A.A. Writing Original Draft: I.L., A.S. Writing Review and Editing of Final Draft: I.L., K.C., A.S., M.M., S.B., V.A., B.L.M, H.A.A., J.L.M. Funding: B.L.M, H.A.A., J.L.M. All authors have read and approved the final submitted manuscript.

## Additional Information

### Ethics Approval

Monocytes were obtained from healthy donors with informed consent under an IRB-approved protocol at the University of Rochester (STUDY0004777). Tenocytes were isolated from tendon tissues obtained during hand surgeries with informed consent under an IRB-approved protocol at the University of Rochester (STUDY00004840).

