## Supplemental Information for "Fluid Flow Impacts Endothelial-Monocyte Interactions in a Model of Vascular Inflammatory Fibrosis"

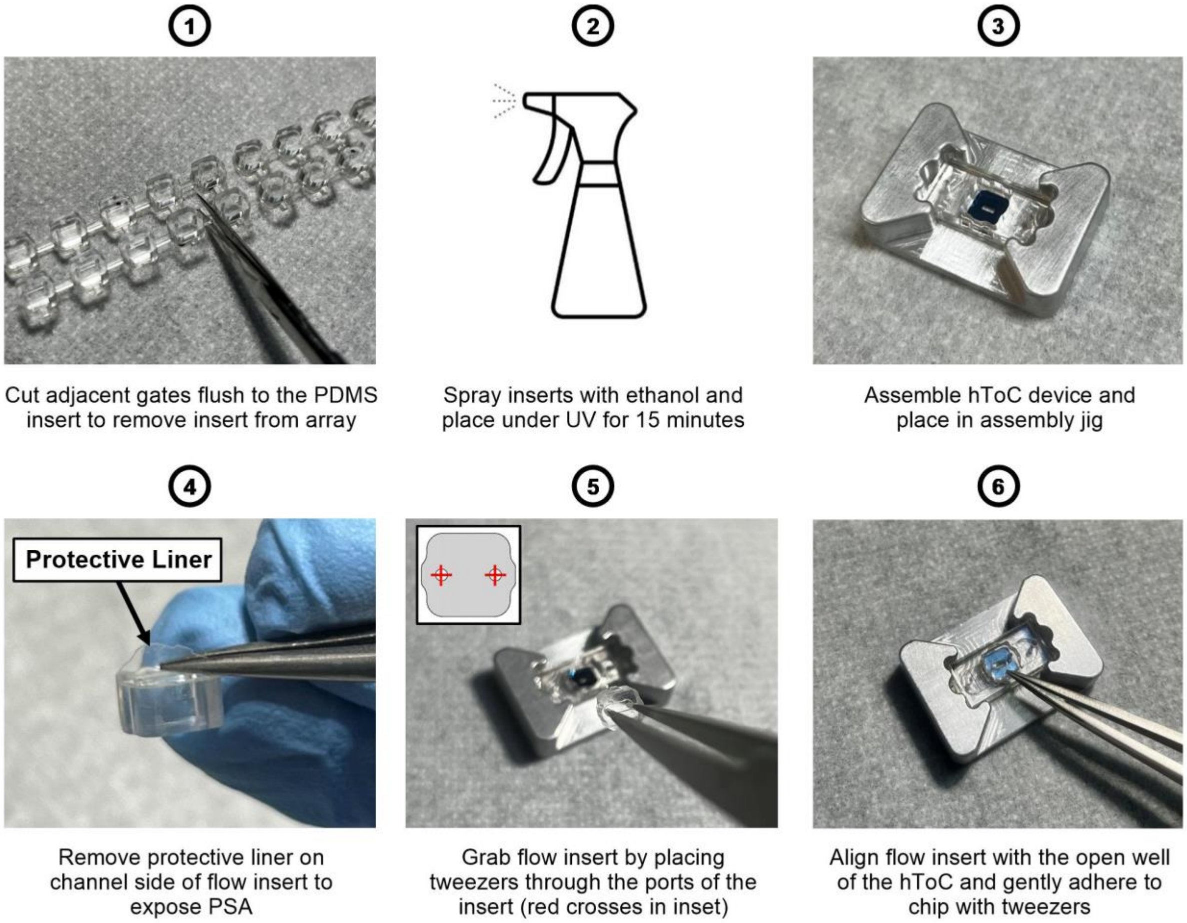


***Figure S1. Assembly steps for manufactured flow insert.*** First, the bridging gates in the manufactured flow insert array are cut out to remove the individual components. The inserts are sprayed with ethanol and then subjected to 15 minutes of UV exposure in the biological safety cabinet. Next, the hToC device is assembled and placed in the assembly jig. The protective liner on the bottom of the flow insert is removed to expose the adhesive, and the insert is placed in the open well of the assembled hToC device. The insert can be gripped by placing the tweezers in the two ports and aligning the insert in the open well of the device. The insert should not be squeezed to avoid deforming the PDMS when placing in the device. The flat side of the tweezers can be used to gently press around the edges of the insert to ensure it is fully adhered to the chip. Avoid pressing down on the middle of the insert, as this may apply pressure to the porous membrane area.

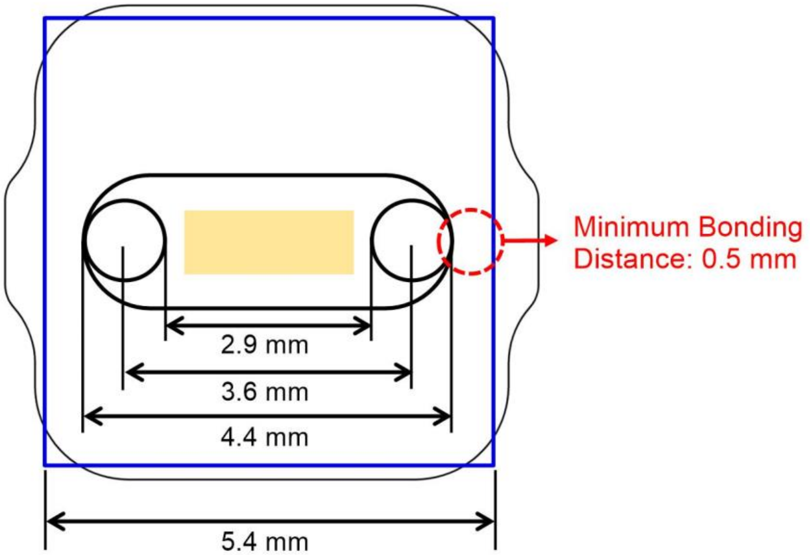


***Figure S2. Cross section of flow insert when adhered to the chip***. Features of the flow insert and key dimensions are outlined in black, the silicon nitride chip is outlined in blue, and the porous membrane area is boxed in yellow. The text in red demonstrates the threshold for minimum bonding distance between the edge of the flow insert and the chip to ensure a leak-proof seal and adhere to manufacturing tolerances.

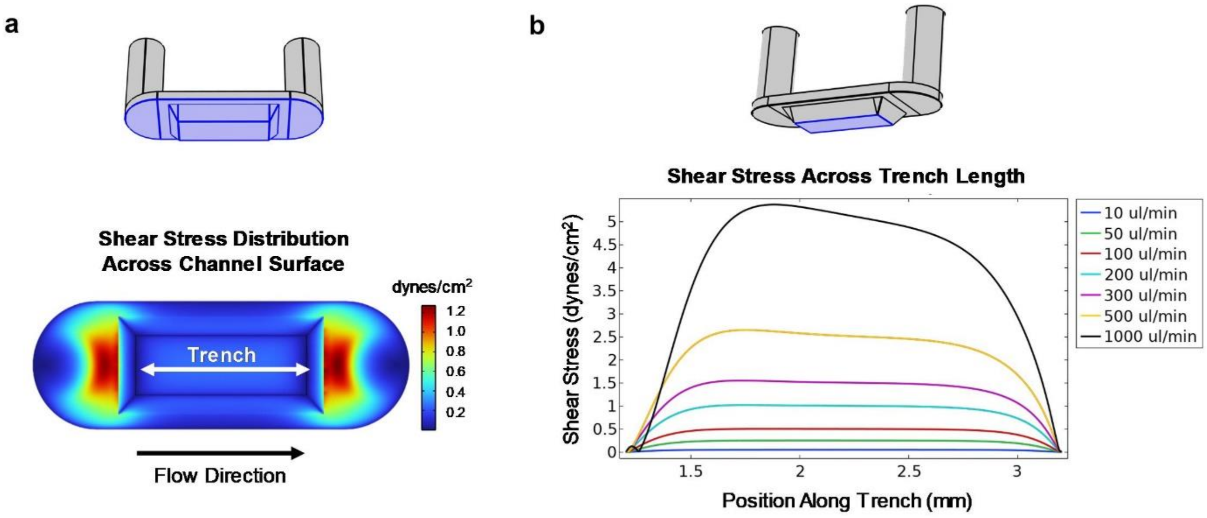


***Figure S3. COMSOL simulation results for 'trench up' membrane configuration.*** **a.** Shear stress distribution across bottom of flow channel with 'trench up' membrane at a flow rate of 50 µl/min, confirming a uniform flow profile across the culture region of the trench. **b.** Plot of shear stress values at the trench surface for flow rates between 10-1000 µl/min.

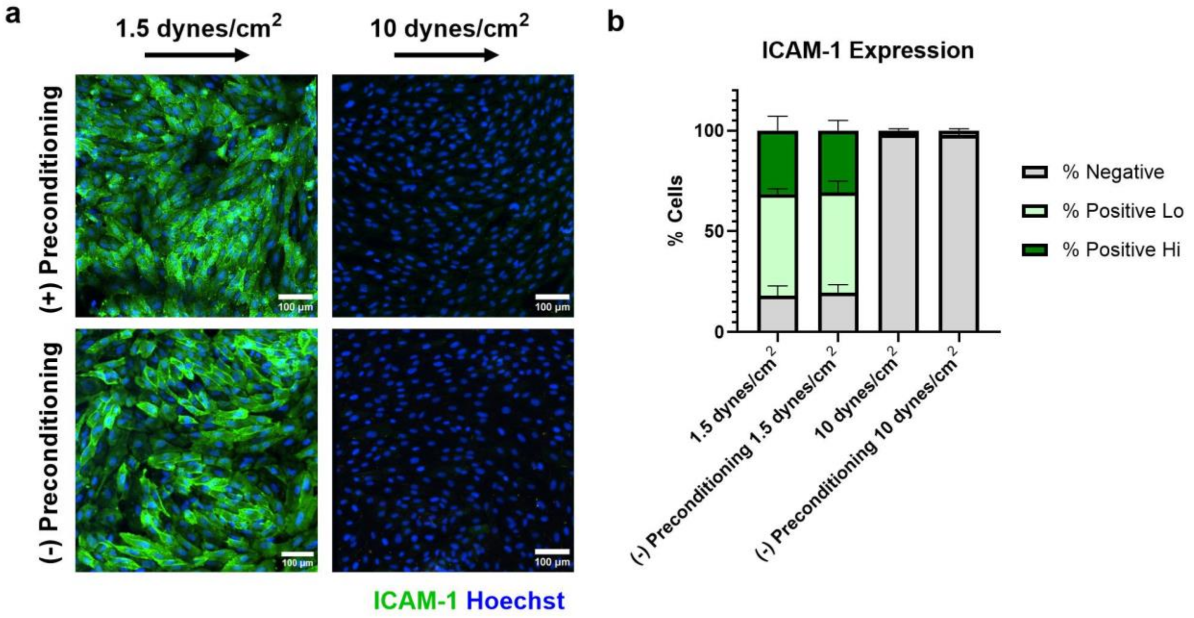


***Figure S4. Comparison of preconditioning versus lack of preconditioning on endothelial ICAM-1 expression with TNF-α stimulation.*** Endothelial cells (ECs) cultured with or without basal 10 ng/ml TNF-α stimulation for 24 hours of fluidic shear stress at 1.5 or 10 dynes/cm$^2$. Preconditioned ECs were subjected to the respective fluidic shear stress for 24 hours prior to TNF-α stimulation. Devices were stained with ICAM-1 (green) and Hoechst (blue). We observed no significant differences between preconditioning and lack of preconditioning on ICAM-1 expression at both low and high shear stresses.

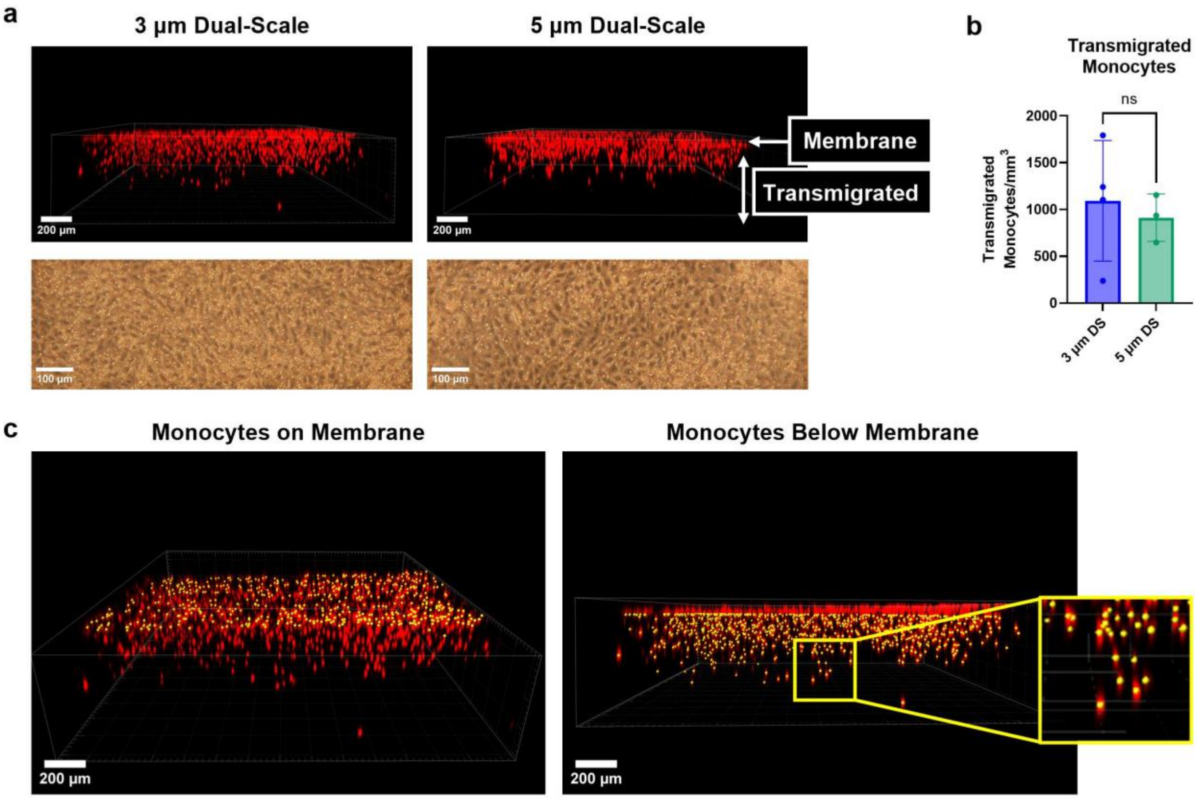


***Figure S5. Comparison of monocyte transmigration across endothelium seeded on 3 or 5 um dual-scale membranes.*** Devices with EC monolayers were preconditioned for 24 hours at 1.5 dynes/cm$^2$ before adding monocytes to the circulation (50 µl/min, 200,000 cells/ml). **a.** Representative confocal images of 3 vs 5 µm devices, with monocytes labeled in red. Phase images show EC monolayers with adhered monocytes after 24 hours of monocyte circulation. **b.** Quantification of transmigrated monocytes at 24 hours of circulation, showing no significant differences in the number of transmigrated monocytes/mm$^3$ between 3 and 5 µm dual-scale devices (unpaired t-test). **c.** Confocal images processed in Imaris demonstrating segmentation of monocytes on or below the membrane. We developed the following image analysis workflow to quantify the monocytes: (1) Identify z slice ranges for monocytes on membrane and transmigrated monocytes, (2) Enter estimated diameter of monocytes, (3) Filter by fluorescence intensity mean, (4) Verify that segmentation parameters have identified all monocytes, (5) Graph quantified values.

***Table S1.*** Key dimensions and volumes of manufactured flow insert

| **Manufactured Flow Insert Metrics** | |
|---|---|
| Channel Height | 0.2 mm |
| Channel Length | 4.4 mm |
| Port Diameter | 0.8 mm |
| Total Channel Volume | 4.23 µl |
| Channel Volume (Minus Ports) | 1.22 µl |

***Table S2.*** Shear stress values from COMSOL simulation comparing ‘trench down’ and ‘trench up’ membrane configurations

| **Inlet Flow Rate (µl/min)** | **Shear Stress on Culture Surface (dynes/cm$^2$)** | | **Max Shear Stress in Channel (dynes/cm$^2$)** | |
|---|---|---|---|---|
| | ***Trench Down*** | ***Trench Up*** | ***Trench Down*** | ***Trench Up*** |
| 10 | 0.18 | 0.05 | 0.4 | 0.25 |
| 50 | 1.5 | 0.25 | 2 | 1.25 |
| 100 | 1.85 | 0.5 | 3.93 | 2.5 |
| 200 | 3.69 | 1.0 | 7.96 | 5 |
| 300 | 5.54 | 1.5 | 11.79 | 3.75 |
| 500 | 9.23 | 2.5 | 20.89 | 12.5 |
| 1000 | 18.45 | 5.0 | 42.49 | 30 |

***Table S3.*** Antibodies for immunofluorescence staining

| **Primary Antibody** | **Dilution** | **Conjugation** | **Manufacturer** | **Catalog #** | **Secondary Antibody** | **Dilution** | **Manufacturer** | **Catalog #** |
|---|---|---|---|---|---|---|---|---|
| **ICAM-1** | 1:100 | N/A | BioLegend | 353102 | Alexa Fluor 488 | 1:200 | Thermo Fisher | A11001 |
| **VCAM-1** | 1:100 | N/A | BD Biosciences | 555645 | Alexa Fluor 488 | 1:200 | Thermo Fisher | A11001 |
| **VE-cadherin** | 1:100 | N/A | R&D Systems | MAB8381 | Alexa Fluor 488 | 1:200 | Thermo Fisher | A11001 |
| **PECAM-1** | 1:100 | N/A | Invitrogen | PA5-32321 | Alexa Fluor 546 | 1:200 | Thermo Fisher | A11056 |
| **PSGL-1** | 1:150 | N/A | R&D Systems | FAB9961G | | | | |
| **Phalloidin** | 1:1000 | iFluor 488 | Abcam | ab176753 | | | | |
| **Hoechst 33342** | 1:2000 | N/A | Invitrogen | H3570 | | | | |

**Supplementary Video Legend**

***Supplementary Video 1.*** Representative video showing the endothelium with circulating monocytes at a flow rate of 50 μl/min and a density of 200,000 cells/ml. Monocyte chemoattractant protein 1 (100 ng/ml) was added to the bottom channel of the device to induce transmigration. The video demonstrates a robust transmigration response with circulating flow, in which phase dark monocytes have transmigrated across the endothelium. Real-time imaging was carried out at 0.125 Hz frame rate.
